# Interest of phenomic prediction as an alternative to genomic prediction in grapevine

**DOI:** 10.1101/2021.12.16.472608

**Authors:** Charlotte Brault, Juliette Lazerges, Agnès Doligez, Miguel Thomas, Martin Ecarnot, Pierre Roumet, Yves Bertrand, Gilles Berger, Thierry Pons, Pierre François, Loïc Le Cunff, Patrice This, Vincent Segura

## Abstract

Phenomic prediction has been defined as an alternative to genomic prediction by using spectra instead of molecular markers. A reflectance spectrum reflects the biochemical composition within a tissue, under genetic determinism. Thus, a relationship matrix built from spectra could potentially capture genetic signal. This new methodology has been successfully applied in several cereal species but little is known so far about its interest in perennial species. Besides, phenomic prediction has only been tested for a restricted set of traits, mainly related to yield or phenology. This study aims at applying phenomic prediction for the first time in grapevine, using spectra collected on two tissues and over two consecutive years, on two populations and for 15 traits. First, we characterized the genetic signal in spectra and under which condition it could be maximized, then phenomic predictive ability was compared to genomic predictive ability. We found that the co-inertia between spectra and genomic data was stable across tissues or years, but variable across populations, with co-inertia around 0.3 and 0.6 for diversity panel and half-diallel populations, respectively. Differences between populations were also observed for predictive ability of phenomic prediction, with an average of 0.27 for the diversity panel and 0.35 for the half-diallel. For both populations, there was a correlation across traits between predictive ability of genomic and phenomic prediction, with a slope around 1 and an intercept of −0.2, thus suggesting that phenomic prediction could be applied for any trait.

## 1 Introduction

Viticulture has to face two major threats in the next decades, diseases and climate change, which impact both yield and wine quality. Plant breeding could help mitigating these impacts by mobilizing grapevine genetic diversity (Morales-Castilla et al., 2020). However, grapevine breeding is currently slow because of the long juvenile period and cumbersomeness of field trials. Genomic prediction (GP), first proposed by Meuwissen et al. (2001) is a promising tool to speed up breeding programs and increase selection accuracy, by using genomic information to predict breeding values of candidates to selection. Even though genotyping costs have decreased drastically during the last decades, they can still be prohibitive when hundreds of selection candidates have to be genotyped. That is why Rincent et al. (2018) proposed to switch from genomic markers to near-infrared spectra (NIRS) measured on plant tissues, in a new concept called phenomic prediction (PP). The relationship matrix based on NIRS is indeed expected to share similarities with the genomic relationship matrix, because a reflectance spectrum is determined by the biochemical composition of the analyzed sample (Beer-Lambert law), which in turn is determined by genetic and environmental factors. As PP uses endophenotypes such as NIRS, it may better account for genotype-by-environment interactions than GP, as well as for non-additive genetic effects. In addition, besides being cheaper, NIR measurements are high-throughput, which is required for screening the large populations typically evaluated in breeding programs. One step further, Robert et al. (2021, in press) proposed a definition of genomic-like omics based (GLOB) prediction, which encompasses both phenomic and other omics-based prediction as in Schrag et al. (2018). GLOB is a particular configuration where NIRS (or other omics) used for model training and prediction come from different environments.

Rincent et al. (2018) found that phenomic predictive ability could be higher than genomic predictive ability with wheat grain NIRS and equivalent predictive ability (PA) with poplar wood NIRS for some traits. In wheat, when predicting across environments, PP was still more accurate than GP for most traits. Other studies, such as Lane et al. (2020) in maize reported PA for PP, but in this study, GP was not implemented for comparison. Krause et al. (2019) applied PP in wheat in a single environment with hyperspectral imaging from different phenological stages, they found higher PA with PP than with GP for most time-points studied. Indeed, this might be explained thanks to GxE interactions, because NIRS on training set (TS) and validation set (VS) were measured in a single environment. Several studies also reported an increase in PA when combining genomic and phenomic matrices in a single prediction model (Cuevas et al., 2019; Galán et al., 2020). Nevertheless, PP is still in its infancy, as it has been mostly applied to cereals with grain and leaves as tissues. Many issues remain, in particular which could be the best way to implement PP in breeding programs. In the case of perennial species, such as grapevine, year effect is known to strongly affect phenotype, and how behaves PP in this context remains to be studied. Also, in the case of woody perennial, wood matter offers another kind of material for collecting spectra which could be complementary to leaves. Rincent et al. (2018) found in wheat that combining NIRS collected on leaves and grains could enhance the PA, but the gain was not systematic. More work is thus required to devise a strategy for implementing PP in breeding programs.

In grapevine, GS has been already implemented and gave promising results on different populations (Brault et al., 2021a,b; Flutre et al., 2020; Fodor et al., 2014). However, so far to our knowledge only one study has evaluated PP in woody perennials (in poplar, Rincent et al. (2018)) and consequently none on grapevine. The aim of this study was to understand under which configuration PP could be implemented in grapevine breeding programs. For that, we first provided a thorough characterization of the genetic signal in spectra. Specifically, we performed a co-inertia analysis (Dolédec and Chessel, 1994) to assess the pairwise relationship between genotyping and NIRS matrices. This methodology was already used in ecology and multi-omics studies but has never been applied in this context (Meng et al., 2014; Min et al., 2019). Then, we compared multiple configurations for performing PP, such as using raw NIRS vs derived BLUPs over a single or two years and over a single or two tissues. Finally, two distinct questions, never addressed before in grapevine, were answered: how do phenomic PA performs compared to genomic PA? Can adding NIRS to genotypic data increase PA?

## 2 Materials and Methods

### 2.1 Plant Material

Our plant material is composed of a diversity panel reflecting the whole genetic diversity of *Vitis vinifera* (Nicolas et al., 2016) and a half-diallel (Tello et al., 2019), better reflecting populations used in breeding programs.

The diversity panel is composed of 279 varieties, with an equal proportion of individuals from each of the three gene pools: Wine West (WW), Wine East (WE) and Table East (TE) (Nicolas et al., 2016). This panel was overgrafted on Marselan in 2009, itself grafted on Fercal. Field location is in the Domaine du Chapitre experimental vineyard of Institut Agro | Montpellier SupAgro in Villeneuve-lès-Maguelone (South of France). The panel is replicated in five randomized complete blocks, each variety being represented by one plot of a single vine in each block.

The half-diallel is composed of 676 individuals from ten bi-parental populations (hereafter named crosses) in a half-diallel mating design between five parents: Syrah (S), Grenache (G), Cabernet-Sauvignon (CS), Terret Noir (TN) and Pinot Noir (PN) (Tello et al., 2019). All of them, except Grenache, belong to the WW gene pool (Brault et al., 2021b). Each cross comprises between 64 and 70 offspring. This population was planted in 2005 and grafted on Richter 110. Field location is the same experimental vineyard, a few kilometers away from the diversity panel field trial. The half-diallel is replicated in two randomized complete blocks, each offspring being represented by one plot of two consecutives vines in each block.

### 2.2 Phenotyping

We studied the same 15 traits in both trials (diversity panel and half-diallel), related to (i) berry composition at harvest, with malic acid (**mal.ripe**), tartaric acid (**tar.ripe**), shikimic acid (**shik.ripe**) concentrations, and shikimic / tar-taric acid (**shiktar.ripe**) and malic / tartaric acid (**maltar.ripe**) ratios, (ii) berry and cluster morphological traits, with mean berry weight (**mbw**), mean cluster weight (**mcw**), mean cluster length (**mcl**), mean cluster width (**mcwi**) and cluster compactness (**clucomp**), (iii) phenology traits, with véraison date (onset of ripening, **verday**), harvest date (**samplday**) and the interval between véraison and harvest (**vermatu**), (iv) vigor (**vigour**). Details about phenotypic measurements, statistical processing and heritability can be found in Brault et al. (2021b). For prediction, we used the Best Linear Unbiased Predictors (BLUP) of genotypic values from Flutre et al. (2020) in the diversity panel and Brault et al. (2021b) in the half-diallel. Briefly, a mixed linear model was fitted for eliminating experimental confounding effects and in order to extract BLUPs of genotypic values. In the following, only BLUPs of genotypic values were used for the diversity panel, whereas the sum of genotypic and cross BLUPs were used for the half-diallel.

### 2.3 SNP genotyping

We used a set of 32,894 SNP markers common to both populations. Details about genotyping and marker processing are given in (Tello et al., 2019) for the half-diallel and in Flutre et al. (2020) for the diversity panel. The selection of common SNPs was done in Brault et al. (2021b). 622 out of 676 individuals were successfully genotyped in the half-diallel, and 277 out of 279 individuals in the diversity panel.

### 2.4 Spectra measurements

Spectra were measured in both trials on dried wood and leaves collected during two consecutive years (2020 and 2021). For wood tissue, two shoots were cut per plot, on two vines in the half-diallel and one in the diversity panel. These wood shoots were approximately 3 cm long. Wood was harvested on January 27^*th*^ in 2020 and January 14^*th*^ in 2021. For leaf tissue, four discs were sampled per plot, on two adult leaves per vine for two different vines in the half-diallel and on four leaves per vine in the diversity panel. Leaf disks had diameters of *circa* 1 cm and 0.5 cm in 2020 and 2021, respectively. Leaf tissue harvest occurred on July 1^*st*^ 2020 and June 16^*th*^ 2021. Two blocks were used in both trials, leading to a total of four wood shoots and eight leaf discs per genotype. After harvest, shoots and leaves were dried at 60°*C* until the weight stopped decreasing, and then stored in a cold chamber until measurements.

For spectra gathering, a reflectance probe plugged to a visible-infrared spectrometer was used (LabSpec 2500 Portable Vis/Nir spectrometer device; Analytical Spectral Devices, Inc., Boulder, CO, US) with its associated software IndicoPro 6.5. A reference spectrum was taken twice a day, using Spectralon ^®^. For each wood shoot, two scans were taken, one on each end of the shoot. For each leaf disc, one scan was taken, on the adaxial surface. Thus, for each tissue, four scans were produced per plot (i.e., per genotype x block combination). Wavelengths ranged from 350 to 2,500 nm, with a 1 nm step. For each scan, the spectrometer takes 10 spectra which are automatically averaged to make one spectrum record. In total around 1,800 and 5,400 scans were collected on the diversity panel and the half-diallel populations for each year and tissue, respectively.

### 2.5 Spectra pre-processing

Spectra were processed separately within each trial. The first 50 wavelengths (visible range) were cut, because of instabilities. The average of the four spectra per plot were then carried out over the 2,101 remaining wavelengths. From these averaged raw spectra (**raw**), five pre-processing were then applied: smoothing (**smooth**) using Savitzky and Golay (1964) procedure, normalization or standard normal variate (**snv**) which consists in centering and scaling (Barnes et al., 1989), detrend (**dt**) for removing baseline (Barnes et al., 1989), and first and second derivative on normalized spectra (**der1**and **der2**, respectively), also for removing baseline and exacerbate some parts of the signal.

On each of these six spectra matrices (**raw**, **smooth**, **snv**, **dt**, **der1**and **der2**), we applied a mixed model over the reflectance at each wavelength, to compute variance components and derive NIRS genotypic BLUPs for each possible combination of three models at the tissue level times three models at the year level (Table 1).

**Table 1:**
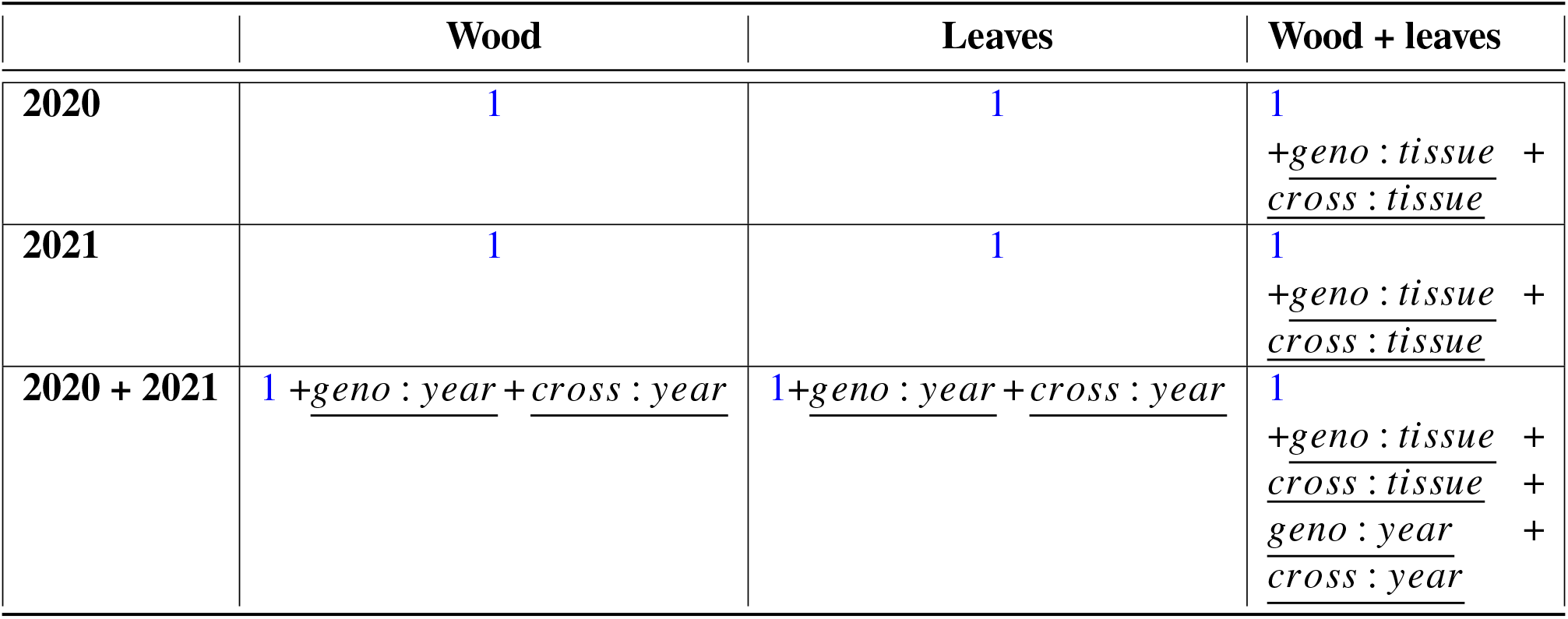
Mixed model fitted, depending on the model combination. *cross* effect is replaced by *subpop* for the diversity panel.

|  | Wood | Leaves | Wood + leaves |
| --- | --- | --- | --- |
| **2020** | 1 | 1 | 1<br>+<u>geno : tissue</u> +<br><u>cross : tissue</u> |
| **2021** | 1 | 1 | 1<br>+<u>geno : tissue</u> +<br><u>cross : tissue</u> |
| **2020 + 2021** | 1 +<u>geno : year</u> + <u>cross : year</u> | 1 +<u>geno : year</u> + <u>cross : year</u> | 1<br>+<u>geno : tissue</u> +<br><u>cross : tissue</u> +<br><u>geno : year</u> +<br><u>cross : year</u> |

The base mixed model was:

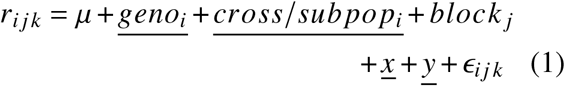

With *r* the reflectance at a given wavelength, *μ* the intercept, *geno* the random genotypic effect, *cross subpop* the random effect for cross (10 levels in the half-diallel) or subpopulation (3 levels in the diversity panel) effect, *block* the fixed effect of block, *x* and *y* the random effects for plot coordinates, and *ϵ* the residuals.

Factors could then be added to this base model, depending on the model combination (Table 1).

NIRS BLUPs used further were the sum of genotypic and cross or subpopulation BLUPs. For comparison, we also computed genotypic BLUPs from models without *cross* or *subpopulation* effects.

For comparison purpose and to evaluate the benefit of fitting a mixed model per wavelength to extract a genotypic BLUP, we also computed for each of the 6 spectra matrices (**raw**, **smooth**, **snv**, **dt**, **der1** and **der2**) the averaged spectra per genotype.

### 2.6 Variance components and co-inertia

Variance components from mixed models were extracted at each wavelength and compared between model combinations and populations.

We also compared relationship matrices obtained independently from SNPs (that is, the genomic relationship matrix, GRM) and NIRS BLUPs (that could be called the phenomic relationship matrix), using co-inertia analysis (Dolédec and Chessel, 1994). Briefly, the co-inertia between two matrices X and Y (from SNP and wood NIRS for example) is computed as:

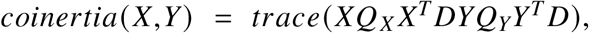

with *Q*_*X*_ and *Q*_*Y*_ the weights associated with X and Y columns (SNP markers and reflectances), which were set to 1, and D the weights associated with X and Y rows (individuals), which were set to 1/n with n the number of individuals.

Then, a measure of correlation between X and Y can be computed as the RV coefficient:

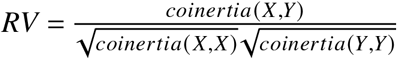

We applied co-inertia analysis to SNPs, wood and leaf NIRS BLUPs, in order to estimate pairwise RV coefficients between these matrices.

### 2.7 Heritability assessment

Heritability values of phenotypic data were assessed for both populations in Flutre et al. (2020) for the diversity panel and in Brault et al. (2021b) for the half-diallel.

Heritability values were also assessed for reflectance data a t each wavelength, a fter mixed model fitting. As for phenotypic data, heritability formula was:

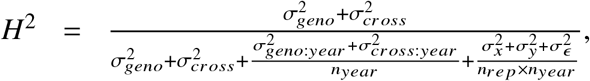

with the variance components previously estimated in the mixed model, *n*_*rep.year*_ the mean number of replicates per year, and *n*_*year*_ the number of year (one or two, depending on the model).

### 2.8 Phenomic and genomic prediction models

Three methods were compared for the implementation of PP and GP, based on two models. Models were fitted separately for each population and trait.

#### 2.8.1 rrBLUP vs GBLUP/HBLUP model type

- In rrBLUP, we fitted the following model: *y* = *Xβ + ϵ*, with *y* the vector of genotypic BLUPs from phenotypic data, *X* the matrix for marker genotypes (additively coded as in Brault et al. (2021b)) or wavelength data (from NIRS BLUPs for each of the nine above-mentioned model combinations), *β* the marker or wavelength effects and *ϵ* the residual effects. This model was fitted using R/glmnet package version 4.1-2 (Friedman et al., 2010). In rrBLUP, marker or wavelength effects are shrunk towards zero, according to a regularization parameter, chosen by an inner cross-validation (CV).
- In HBLUP (GBLUP model, using the NIRS relationship matrix H), we fitted the following model: *y* = *u* + *ϵ*, with *y* the vector of genotypic BLUPs from phenotypic data, *u* the random effects for genomic or phenomic estimated breeding value, with 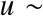 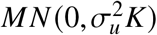, *K* being the relationship matrix from markers or spectra, 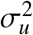 the genetic variance, and *ϵ* the random residual effect, 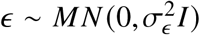, *I* being the identity matrix. 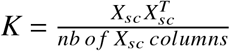, *X*_*sc*_ described X matrix scaled on allelic frequencies or wavelength reflectances. This model was fitted using R/lme4GS package version 0.1 (Caamal-Pat et al., 2021).

#### 2.8.2 Cross-validation

PP and GP models were assessed within each population and for each trait using CV. In the half-diallel, 10-fold CV was applied, while in the diversity panel, 5-fold CV was applied; CV was repeated 10 times. For each CV replicate, predicted values from all folds were combined and compared with observed genotypic BLUPs. We computed predictive ability (PA) as Pearson’s correlation between the observed and predicted genotypic values. In the half-diallel, PA was calculated within each cross, as it was done in Brault et al. (2021b).

#### 2.8.3 Multi-matrix model fitting

Using R/lme4GS allowed us to fit a single model involving several variance-covariance matrices, such as: 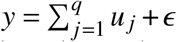, with 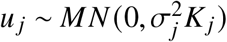, and *K*_*j*_ the relationship matrix from SNPs, wood NIRS or leaf NIRS previously described. We fitted this multi-matrix model using two or three variance-covariance matrices: SNPs + wood NIRS, SNPs + leaf NIRS, wood NIRS + leaf NIRS and SNPs + wood NIRS + leaf NIRS.

## 3 Results

### 3.1 Characterization of genetic signal in spectra

#### 3.1.1 Variance components

Variance components for the nine model combinations studied in each population are shown in Figure 1 for **der1** pre-processing. In both populations, genotypic variance was maximized in single-year and single-tissue analyses, with an essentially comparable magnitude between wood and leaves on the one hand and 2020 and 2021 on the other hand. In multi-tissue analyses, genotypic variance drastically decreased and was mostly replaced by *geno:tissue* variance, while in multi-year analyses, genotypic variance was only partly replaced by *geno:year* variance. A strong *x* effect (row effect) was observed, while barely no *y* effect was present.

**Figure 1:**
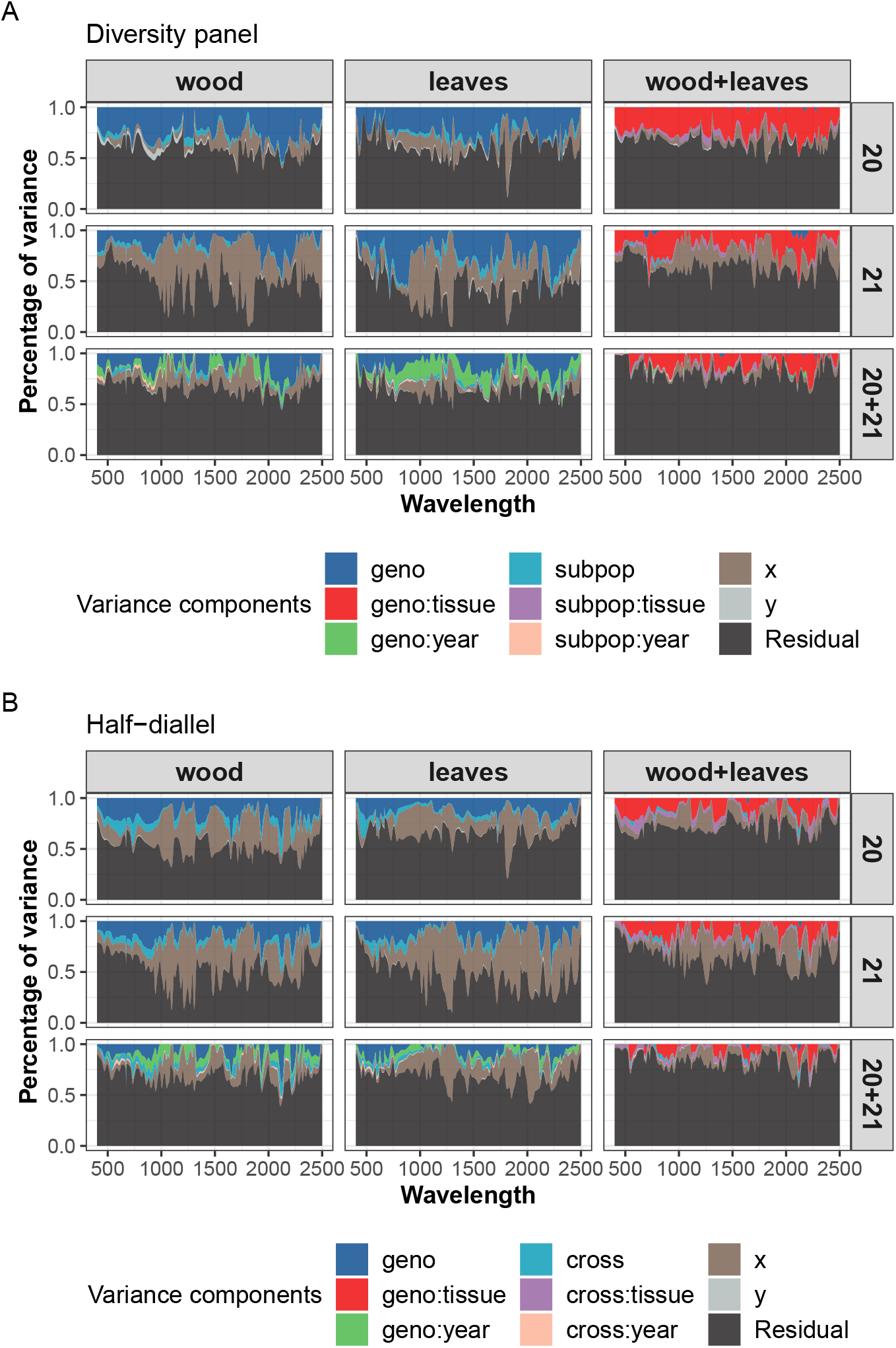
Variance components from the mixed models applied to NIRS after **der1** pre-process. A: in the diversity panel population, B: in the half-diallel population.

Comparing populations, the *cross* variance in the half-diallel was larger than the *subpop* variance in the diversity panel. The variance of interactions between *cross* or *subpop* and *year* or *tissue* remained low. The *geno:year* interaction was more important in the diversity panel than in the half-diallel.

#### 3.1.2 Heritability

Genotypic variance results were consistent with heritability values calculated for each wavelength (distributions of heritability values for each pre-process are given in Figure S1).

When comparing raw and pre-processed spectra, it was clear that the lowest heritability values generally corresponded to **raw** and **smooth** spectra. Heritability values for other pre-processes were close to each other, **der1** yielding the highest heritability values overall (Figure S1).

Including both wood and leaf NIRS in the mixed model resulted in very low heritability values (Figure S1), hence we excluded this model in the following analyses. The analysis wavelength by wavelength has showed that NIRS carry some genetic variance, with a moderate magnitude. To further characterize this genetic signal over the entire spectral range, we then carried out a co-inertia analysis between NIRS and SNP matrices.

#### 3.1.3 Comparison of matrices from SNPs and NIRS, using co-inertia analysis

Co-inertia analysis was conducted on single-tissue models only. Figure 2 shows for each population the relative co-inertia between three matrices of SNPs, wood and leaf NIRS BLUPs of *genotype + cross* or subpopulation effects for “2 years” models. For both populations, correlation with SNPs was similar between wood and leaf NIRS. However, this correlation was nearly twice higher in the half-diallel than in the diversity panel. It is noteworthy that in both populations the correlation between the SNP matrix and NIRS BLUPs matrices (obtained from wood or leaves) was higher than between the two NIRS BLUPs matrices obtained on wood and leaves.

**Figure 2:**
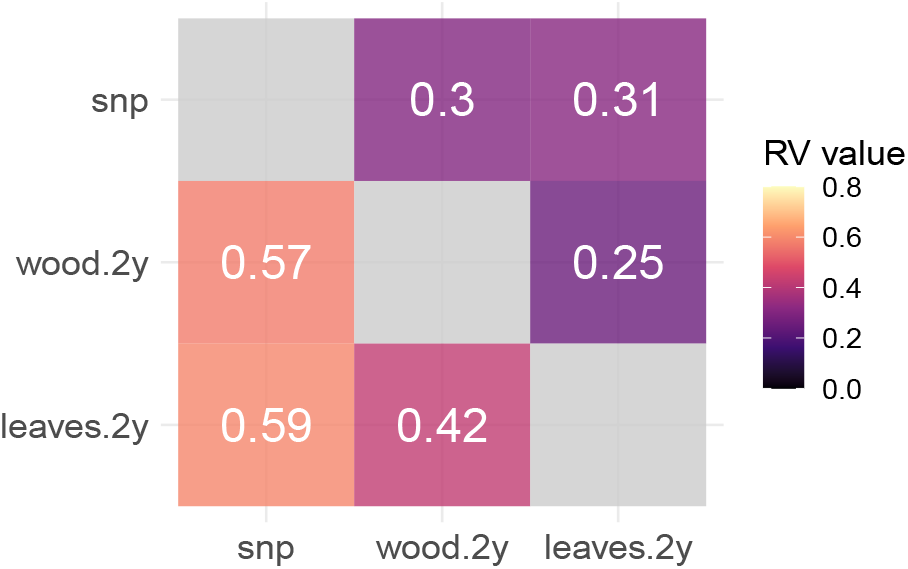
Correlation between the genomic relationship matrix (“snp”) and the relationship matrices derived from wood and leaf NIRS BLUPs of genotype + cross or subpopulation effects with both years included in the mixed model (“wood.2y”, “leaves.2y”, respectively). Upper triangle: in the diversity panel, lower triangle: in the half-diallel.

We also carried out the co-inertia analysis with matrices derived from NIRS BLUPs of *genotype* effect for a model containing either a *genotype* effect only or both *genotype* and *cross* or *subpopulation* effects (Figure S2).

Using such matrices strongly decreased correlation with the SNP matrix, as compared to using matrices derived from BLUPs of *genotype + cross* or *subpopulation* effects (Figure S2). Therefore, in subsequent prediction analyses, we used only the latter matrices including *cross* or *sub-population* effect. Matrices from multi-year NIRS BLUPs generally displayed a slightly higher correlation with the SNP matrix than the single-year BLUPs, and this effect was more pronounced in the half-diallel (Figure S3).

### 3.2 Phenomic prediction using BLUPs vs *base* spectra and rrBLUP vs HBLUP

In each population, using spectra BLUPs instead of *base* spectra almost always resulted in higher PA (Figure 3). However, differences were observed depending on the method and population. The method yielding the highest PA was HBLUP (implemented with lme4GS) in the half-diallel and rrBLUP (implemented with glmnet) in the diversity panel. However, it is worth mentioning that differences between methods were found to be more pronounced in the half-diallel than in the diversity panel. The highest differences between *base* spectra and BLUPs were observed for the best method in each population. Thus, we retained spectra BLUPs in all cases, lme4GS in the half-diallel and glmnet in the diversity panel.

**Figure 3:**
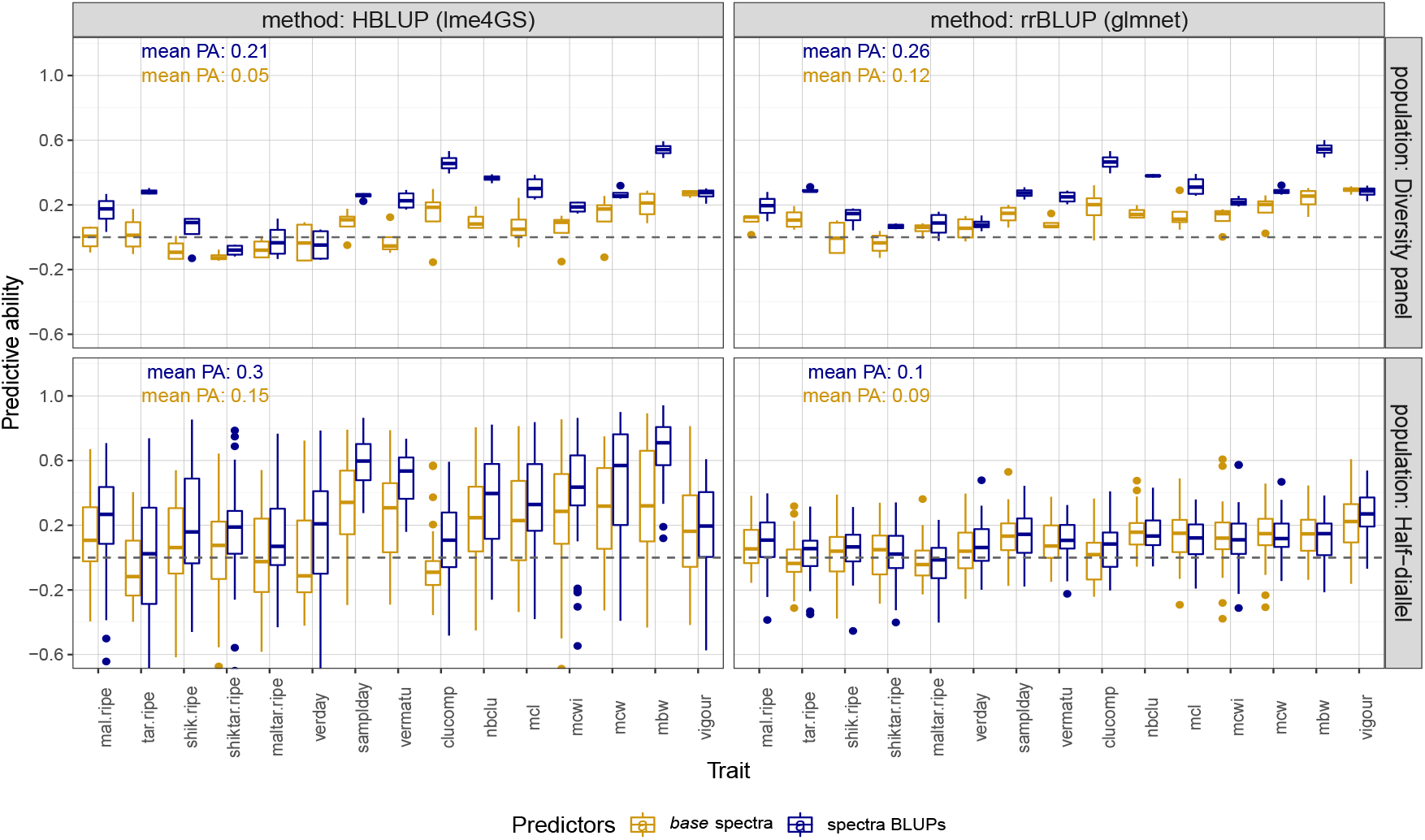
Predictive ability of phenomic prediction for 15 traits, in two populations with two methods, using *base* spectra (in golden) or spectra BLUPs (in dark blue) after der1 pre-process. For each trait, PA distribution was over the 6 models retained for years and tissues (and also over the 10 crosses in the half-diallel).

We observed higher variance in PA in the half-diallel than in the diversity panel, because in the half-diallel, PA distribution was over 10 crosses in addition to the 6 years x tissues model combinations retained (see above). Average PA for the best method was slightly higher in the half-diallel (0.3) than in the diversity panel (0.26).

We compared PA for all pre-processes, after selecting the best method for each population (Figure S4). We found that **der1** and **der2** pre-processes gave close results, with a slight superiority of **der1** overall. Therefore, we kept only this pre-process in subsequent analyses.

### 3.3 Phenomic prediction using NIRS collected over one or two years and tissues

We further compared PP models including a single vs both NIRS BLUP matrices obtained from wood and leaves. For each tissue, we used the NIRS BLUPs derived from the above-described years (2020, 2021 or both). For single tissues, we used the best method selected above in each population, and one NIRS BLUP matrix was fitted. For the wood+leaves configuration in both populations, two NIRS BLUP matrices (one for wood and one for leaves) were fitted using lme4GS package.

For both populations, the nine configurations tested resulted in close PA distributions (Figure 4). Yet, “two years” and “two tissues” configurations overall gave the best average PA values. We thus retained only multi-year and multi-tissue PP results for subsequent comparison with GP.

**Figure 4:**
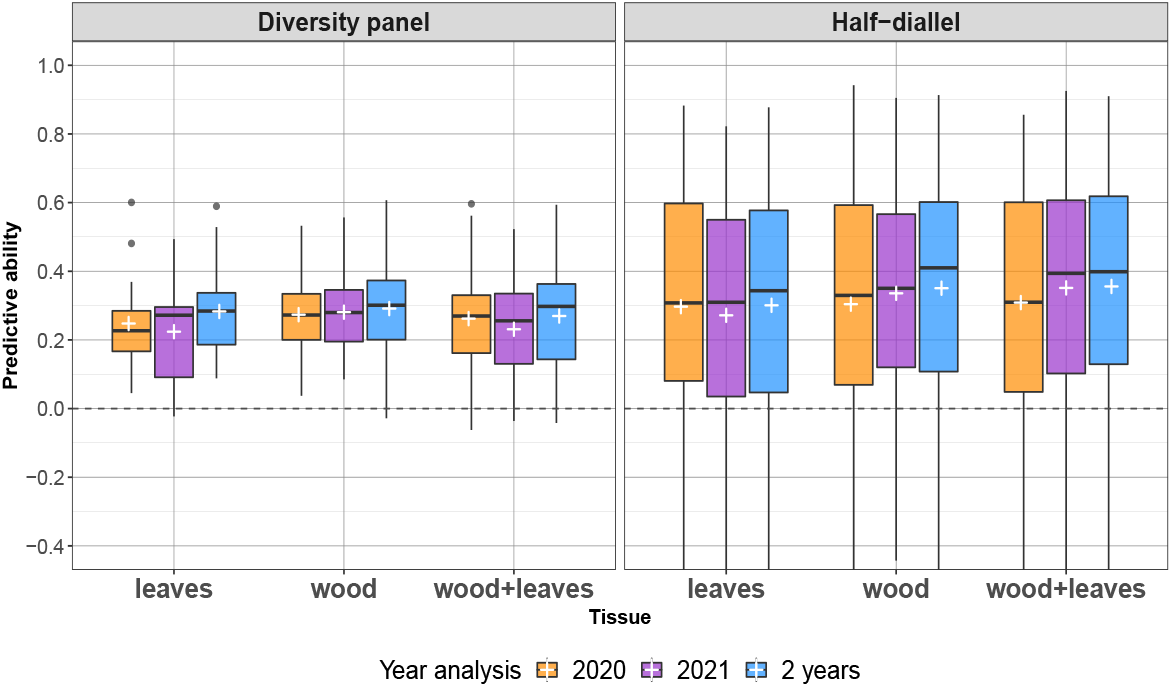
Predictive ability of phenomic prediction with a single vs both years and tissues, over the 15 traits in both populations and the 10 crosses in the half-diallel. Prediction models were fitted with glmnet in the diversity panel (except for wood+leaves configuration) and with lme4GS in the half-diallel. In both populations, models were carried out after der1 pre-processing. The white cross indicates the average PA for each configuration.

Finally, PA was slightly higher in the half-diallel on average, with a larger variance originating from differences between crosses (see hereafter). If we now turn to details per trait, the results show that, within the diversity panel, each trait displayed nearly the same PA for the different tissues, for PA values above 0.2 (Figure S5). In the half-diallel, there were larger differences between tissues. However, this factor still had far less impact on PA than cross or trait.

Overall average PA of PP for “2 years” and “wood+leaves” was 0.27 in the diversity panel and 0.36 in the half-diallel (Figure 4). PA values per trait ranged from −0.04 for **shiktar.ripe** to 0.59 for **mbw** in the diversity panel (Figure S5A), and from 0.13 for **tar.ripe** to 0.74 for **mbw** in the half-diallel (note that in the half diallel, PA values per trait are averaged over the 10 crosses). However, large differences in PA of PP were observed within a trait at the cross level in the half-diallel for “2 years” and “wood+leaves” configuration, such as for **mal.ripe**, from −0.56 for TNxG to 0.62 for TNxCS (Figure 5B and Figure S5B). Comparatively, differences at the cross level were lower for GP (Figure S6). The best predicted cross with PP over all traits was GxS (average PA of 0.41) and the worst one was TNxG (0.28) (Figure S7). For some crosses and traits, PA values could be above 0.8, the maximum PA for PP being 0.91 for **mbw** and TNxG (Figure S5B).

**Figure 5:**
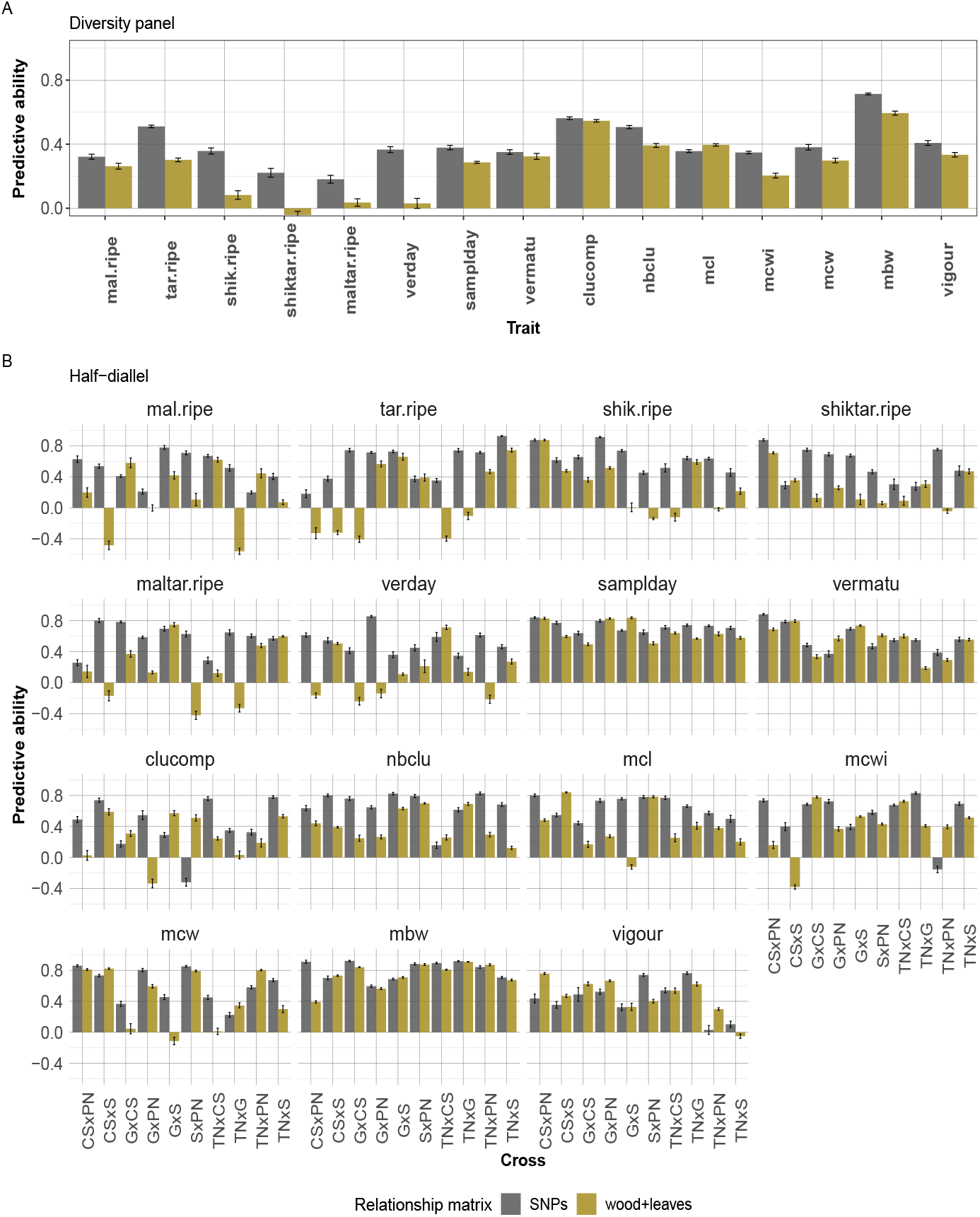
Predictive ability in two settings for 15 traits with lme4GS: SNPs: GP; wood+leaves: PP with two variance-covariance matrices, for wood and leaf NIRS, for “2 years” NIRS BLUPs derived after der1 pre-process. Prediction models were fitted with lme4GS. Error bars correspond to the 95% confidence interval around the mean, based on CV repetitions.

### 3.4 Comparison of PP with GP

Before comparing PP with GP, we applied GP on both populations with the two methods previously compared for PP (Figure S6). We found that lme4GS was overall the best method in both populations, hence we retained this method for the following comparison. Like this was the case for PP, differences between methods appeared to be more pronounced in the half-diallel than in the diversity panel.

The PA reached by PP was close to that of GP in both populations for a few traits and for some half-diallel crosses (**samplday**, **vermatu** and **mbw**) (Figure 5). PP even outperformed GP for some crosses and traits in the half-diallel, such as for CSxPN, GxCS, GxPN and **vigour**, or GxCS, SxPN, TNxPN and **clucomp**. Differences in PA between PP and GP were lower in the diversity panel than in the half-diallel.

In the diversity panel, PA of PP was significantly higher (non-overlapping error bar) than PA of GP for one trait (**mcl**) and non-significantly different for two other traits (**clucomp** and **vermatu**) (Figure 5A). In the half-diallel, PA of PP was significantly higher than PA of GP for 28 trait x cross combinations out of 150, while this difference was not significant in 17 other cases (Figure 5B). In all other cases, PA of PP was lower than PA of GP.

In Figure 6, we further compared mean PAs of PP and GP per trait in each population. In both populations, the slope of the regression model was close to 1 and the intercept to −0.2. This suggests that PA of PP and GP follow the same ranking, independently of the trait. However, this regression had a much lower *R*^2^ in the half-diallel than in the diversity panel.

**Figure 6:**
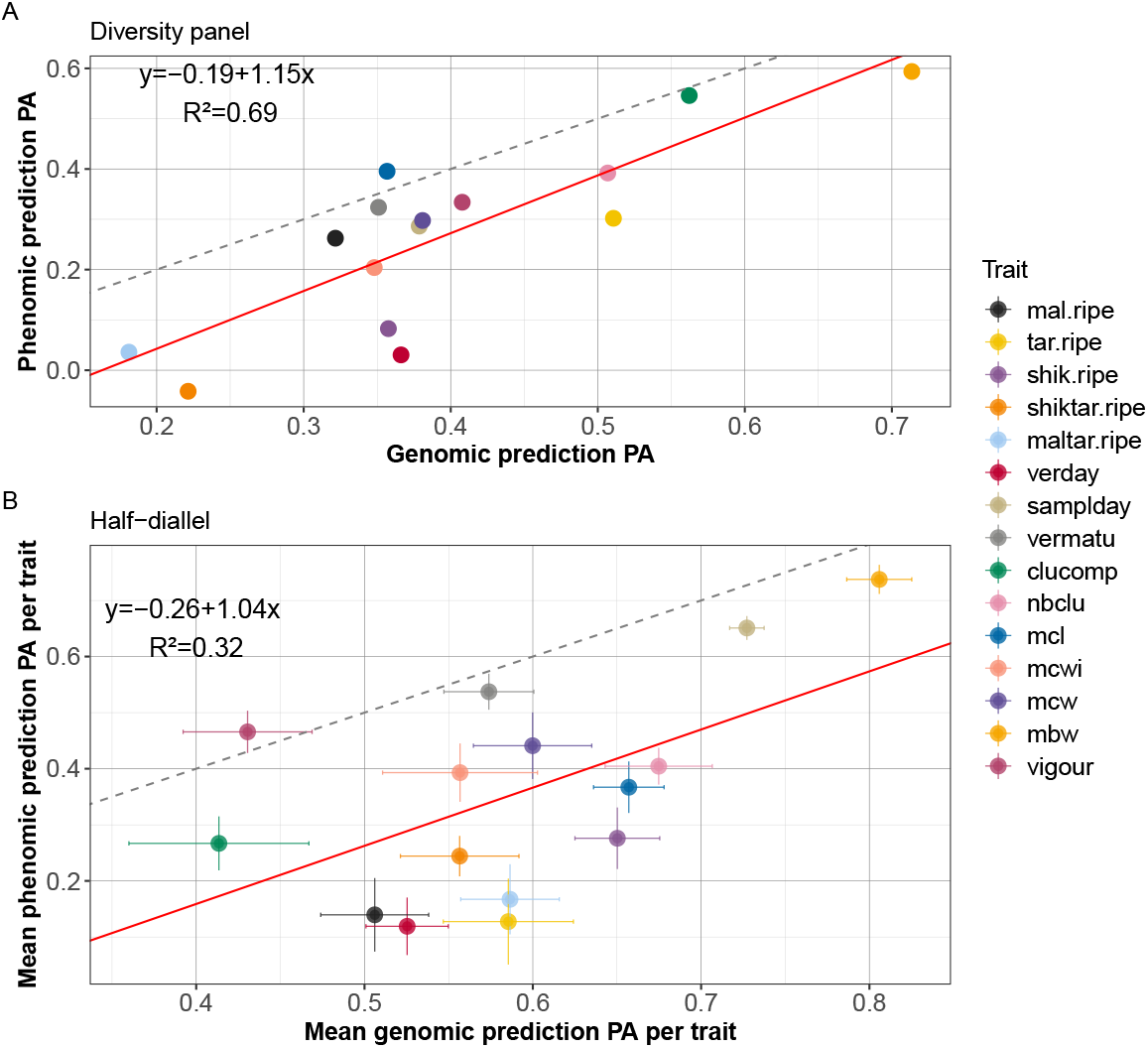
Predictive ability of phenomic against genomic prediction. A: in the diversity panel, B: in the half-diallel. The regression formula corresponds to the following linear model: PA of *PP ~ PA* of GP, with 15 observations (traits). In the half-diallel, PA was averaged across the ten crosses, hence standard error around each point is displayed. The red line is the regression line and the gray dashed line is the identity line.

### 3.5 Enhancing genomic prediction using NIRS

Another possible way of using NIRS is to add it into the predictive model together with SNPs, in order to increase PA. We thus implemented multi-BLUP models with SNPs and NIRS BLUPs and compared them to GP models in each population.

Overall, for both populations and for all traits, differences in PA between SNP based model and different combined GP+PP models were small (Figure S8). In the diversity panel, combining wood NIRS with SNPs led to the best PA (0.406), closely followed by leaves NIRS + SNPs (0.404), wood NIRS + leaves NIRS + SNPs (0.402) and SNPs alone (0.400). In the half-diallel, SNPs alone gave the highest PA (0.590), followed by wood NIRS + leaves NIRS + SNPs (0.564), wood NIRS + SNPs (0.564), and leaves NIRS + SNPs (0.561).

Nevertheless, adding NIRS to a predictive model could lead to minor (non-significant) improvements in PA for some traits, compared to classic GP. Combining GP + PP from wood NIRS slightly increased PA over the GP model for two traits in the diversity panel (**clucomp** and **mcl**) (Figure S8A). In the half-diallel, the difference in average PA with GP was much more variable among traits, with an increase for **vigour**, **clu-comp**, **vermatu**and **samplday**, and a decrease for **mal.ripe**, **tar.ripe**, **shik.ripe**, **shiktar.ripe**, **maltar.ripe**, **nbclu**, **mcl** and **mcwi**(Figure S8B).

## 4 Discussion

So far, PP has only been implemented in a reduced number of species and traits. This study provides the first use of PP in grapevine, within two complementary populations: a diversity panel and a half-diallel. Besides, we tested PP for 15 traits, belonging to four categories: berry composition, phenology, morphological traits and vigour. We first showed that NIRS variability was partly of genotypic origin. We then tested several parameters (mean vs BLUP, tissue, year, method), to optimize both PP and GP. Finally, we found that PP could yield PA values close to or even higher than GP ones.

### 4.1 NIRS variance components and co-inertia with SNPs

Genotype and derived interaction variables had a fairly moderate impact on total variance observed between spectra (Figure 1). The genotypic effect was best captured in single-tissue analyses. This was not surprising, because the genetic signal at a given wavelength relies on molecules specific to each tissue. Then, mixing both tissues into a single model led to no overall genetic effect and to strong *geno:tissue* interaction. This tendency was also observed, to a smaller extent, in the multi-year analyses. This also suggests that different tissues bring non redundant genetic information. This was confirmed by co-inertia analysis, which evidenced that NIRS matrices from wood and leaves were more correlated to the SNP matrix than to each other.

Interestingly, co-inertia analysis showed that multi-year NIRS BLUP matrices were slightly more correlated with the GRM than single-year ones, despite lower genotypic variance. This implies that the genotypic part of NIRS estimated by multi-year analysis could be more related to the genetic signal. Thus, genetic signal ignoring genotype-by-environment interactions could be better captured when several years are combined, this was also the case in Galán et al. (2021) for which multi-year spectra resulted in higher PA values.

Comparatively to genotype-related effects, among non-residual variance components, “x” effect displayed a large variance along wavelengths (Figure 1). This effect actually corresponds to a row effect and might be due to the experimental design. Indeed, leaf discs and wood shoots were both sampled and scanned row by row. However, we cannot determine whether this “x” effect comes from the tissue sampling, i.e., sampling time (over a day), soil heterogeneity; or from the NIRS measurement step, i.e., device calibration, differential storage time, air humidity. Our results underline the importance of accounting such experimental effects in order to improve the genetic signal capture and thus prediction. In further experiments, one could increase the number of spectra per plot and randomize NIRS measurements, in order to determine if the "x” effect observed here was due to measurement or sampling and to reduce it. Other studies that fitted a linear model for each wavelength did not introduce field coordinates as effects (e.g. (Galán et al., 2020; Krause et al., 2019; Lane et al., 2020). But the first and last studies were based on hyperspectral images taken with aircraft flights, that is with an experimental design less prone to plot location effect, and the second study fitted a linear model with only block and environmental effects.

Galán et al. (2020) found a mean heritability value of 0.73 for wavelength reflectances, which is substantially higher than the values we observed (Figure S1). However, we did not use the same heritability formula. Montesinos-López et al. (2017b) also reported overall higher heritability values ranging from 0.6 to 0.8 for most time points, with strong variations depending on the environment (water availability) and time-point.

We found higher heritability and genetic variance in the diversity panel than in the half-diallel. Yet, PA were generally higher in the half-diallel. In Rincent et al. (2018), genetic variance estimates per wavelength between wheat and poplar were consistent with PA in these species, i.e., they evidenced higher PA values in wheat than in poplar. On the opposite, our results on co-inertia analysis were consistent with PA values: correlation between SNP and NIRS BLUPs matrices was higher in the half-diallel than in the diversity panel (Figure 2). This suggest that co-inertia analysis is more relevant to compare configurations for NIRS BLUP than variance components. The higher correlation observed between SNP and NIRS BLUP in the diallel with respect to the diversity panel is likely to be explained by the higher genetic structure in the half-diallel, or because the half-diallel is in better health than the diversity panel, which is older and overgrafted. Actually, it was surprising that NIRS could capture genetic structure, i.e., in our case the *subpopulation* effect in the diversity panel and the *cross* effect in the half-diallel. Although variance components for *subpopulation* and *cross* remained moderate (Figure 1), adding the corresponding BLUP effects to genotypic effects led to a sharp increase in correlation between NIRS and SNP matrices (Figure S2). Further in-depth studies are required to better understand whether this observation could be specific to some subpopulations or families.

### 4.2 Optimizing PP

Among the parameters tested, some had substantial impact on PA, while others had only negligible impact. Namely, using NIRS via BLUP analysis instead of merely average spectra per genotype led to a strong increase in PA (Figure 3). This was probably associated with the strong *x* effect we observed in variance analysis. Such a difference had never been reported before, as studies obtained PP results either from *base* (such as Cuevas et al. (2019); Rincent et al. (2018)) or BLUE (such as Krause et al. (2019); Lane et al. (2020)) spectra, without comparing both configurations.

Surprisingly, the prediction method also had notable impact on PA: using rrBLUP or HBLUP/GBLUP models gave different PA in the half-diallel, while differences in PA between methods were lower in the diversity panel (Figure 3). Yet, HBLUP/GBLUP and rrBLUP models are expected to perform similarly when the regularization parameter in ridge regression is equal to 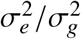 (**?**). In our analysis, this parameter value was chosen by cross-validation using cv.glmnet function. The higher relatedness between genotypes within the half-diallel than within the diversity panel (Brault et al. (2021b), Figure 1a) may boost HBLUP and GBLUP models compared to rrBLUP in this population. In future investigations, one could use variable selection method such as LASSO to select the most relevant wavelengths for computing the relationship matrix from NIRS BLUP. Such variable selection was performed by Galán et al. (2020) and resulted in higher PA.

On the opposite, using single-year, single tissue, multi-year, or multi-tissue NIRS BLUPs and all pre-processes except **smooth** gave very similar results over all traits and crosses (Figure 4), with a slight superiority of multi-year model overall. This was consistent with the results of co-inertia analysis (Figure S3). In Rincent et al. (2018), the multi-tissue analysis for wheat with leaf and grain combined gave similar PA as for single-tissue analysis. As the combination of two tissues for PP was only done in one other study (Rincent et al., 2018), further work needs to be done to assess these conclusions.

For a given trait, both tissues tested gave similar PA for the diversity panel (Figure S5A). For the half-diallel, more differences were observed between tissues, and much larger differences were observed between crosses (Figure S5B). However, no cross was consistently well or poorly predicted for all traits, suggesting a strong cross x trait interaction. These large disparities among crosses were consistent with the GP results obtained in the same population by Brault et al. (2021b).

### 4.3 Comparison between PP and GP

PP is supposed to better account for GxE than GP. However, it was shown in Rincent et al. (2018) that PP could still reach good PA values when NIRS for TS were taken in an environment different from the one in which VS was phenotyped, i.e., when accounting for GxE was not possible. In this study, we could not assess whether PP accuracy partly relied on location-related GxE, because phenotypes and NIRS came from a single location. Nevertheless, phenotypes were measured in 2011-2012 and 2013-2017 in the diversity panel and half-diallel populations, respectively, whereas NIRS were measured in 2020-2021 in both populations. Vintage (year) effect is also part of GxE and it is likely that 2020 or 2021 could display some differences in terms of weather with phenotyping years. For training and validation model, we used genotypic BLUPs of both phenotypic data, thereby removing year and *geno:year* effects. We found that PA seemed not to be impacted by NIRS year for all traits studied, suggesting that vintage has a negligible effect on PA when genotypic BLUPs are used.

As a prospect, one could specifically extract genotype x year and genotype x location variance components from phenotypic and NIRS data and test if PP could be useful to predict this GxE part. Montesinos-López et al. (2017a) studied GxE using spectra and they compared models including or not GxE and wavelength-by-environment interaction. They reported that the inclusion of GxE provided no increase of PA while including wavelength-by-environment interaction was the best model.

We found that PP could compete with GP for some traits in both populations, despite moderate genetic variance estimated from NIRS. However, the number of traits for which PP outperformed GP remained low. These results were close to those of Rincent et al. (2018) on poplar. In our case, one explanation could be that NIRS came from tissues sampled in 2020 and 2021, while phenotypes were measured in 2011-2012 and 2013-2017 in the diversity panel and half-diallel, respectively. Thus, we couldn’t take into account for GxE from vintage effect. As a perspective, it would be interesting to compare PA when spectra are measured the same year as phenotyping or not. In such case, one could explicitly model vintage effects in spectra to further increase PA.

Nevertheless, even when PP does not outperform GP, it may still be interesting in breeding, because of its lower cost and increased throughput compared to genotyping. Moreover, when a trait was well-predicted with GP, we found that it was also well-predicted with PP, with a global shift of −0.2 in PA (Figure 6). This suggests that PP PA truly relies on genetic variability and that PP could be applied indifferently for all traits. Even though this study is the first one implementing PP on so many traits (15), these conclusions remain to be confirmed on other species and traits. Based on data from Rincent et al. (2018) and on the relative GP and PP reliability that we observed, we are still expecting a positive genetic gain by switching from GP to PP.

We tested whether combining NIRS and SNP could increase PA compared to GP, by taking other genetic effects into account. However, as we used NIRS BLUPs, we only maximized the genetic variance part of spectra, we thus intentionally excluded GxE. Therefore, the fact that adding NIRS to GP model did not result in any increase in PA is consistent with our spectra processing. Cuevas et al. (2019) and Galán et al. (2020) found slight to noticeable improvement in PA when NIRS was added to the model, compared to GP model with SNPs only; difference in PA was at most 0.01 in Cuevas et al. (2019) and up to 0.1 in Galán et al. (2020). Both studies are however so different than ours that it is difficult to explain these different behaviors.

As a conclusion, we provided the first implementation of PP in grapevine. The number of traits studied allowed us to put forward a correlation between PA of GP and PP, suggesting that PP relies on a genetic basis. Such a correlation was never reported before. We expect that the shift of PA between PP and GP of −0.2 would be reduced if year of phenotyping and spectra measurement are the same. Still, PP has shown its interest for breeding over a wide range of traits.

## Data availability

All analyses were conducted using free and open-source software, mostly R. Genotypic values and genotypic data for half-diallel and diversity panel populations are available at https://doi.org/10.15454/PNQQUQ. Spectra, R scripts and result tables have been deposited in the INRAE data portal : https://doi.org/10.15454/BICRFX.

## Declarations

### Ethical standards

The authors declare that the experiments comply with the current laws of the country in which they were carried out.

### Author contribution statement

VS, AD, PT and LLC conceived the idea of the study and contributed to funding acquisition; TP, PF and AD obtained the phenotypic data used in this work; YB, GB, MT, JL, VS, LLC and CB measured spectra; ME and PR gave advices on spectrometer use; CB and JL analyzed data and interpreted results with inputs from AD, LLC, PT and VS; PT is the PhD supervisor of CB; CB wrote the original draft, which was reviewed and edited by all authors. All authors read and approved the final manuscript.

## Acknowledgements

This work has been realized with MESO@LR-Platform at the University of Montpellier.

