## Supplemental information for "Interest of phenomic prediction as an alternative to genomic prediction in grapevine"

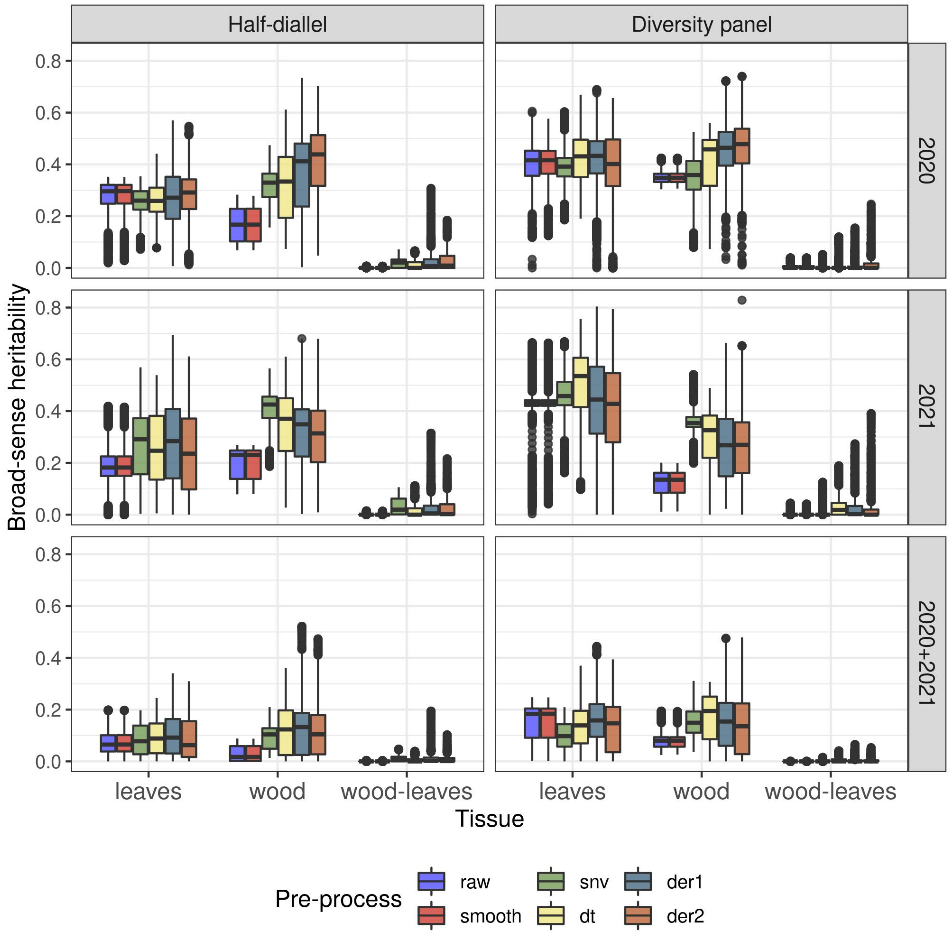


#### Figure S1 Heritability distribution across wavelengths for each pre-process

Comparison of broad-sense heritability between the six NIRS pre-processes, displayed for both populations, three year and three tissue BLUP models.

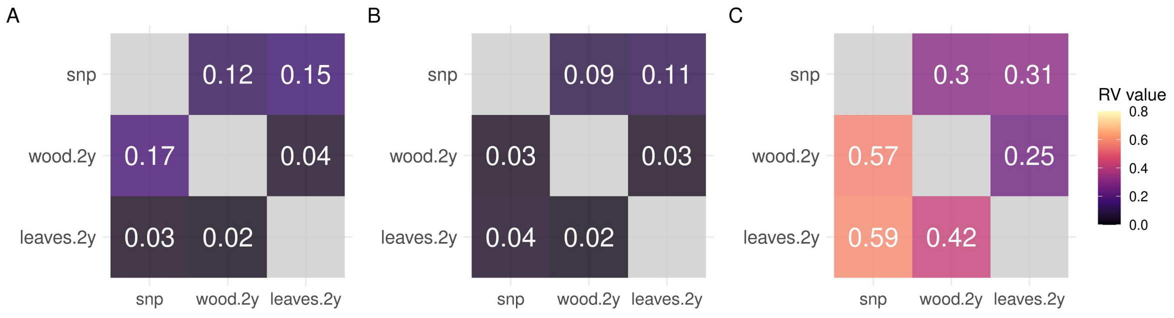


#### Figure S2 Co-inertia analysis between SNP and multi-year wood and leaves NIRS matrices

Upper triangle: in the diversity panel, lower triangle: in the half-diallel

A: mixed model with only genotype effect, resulting NIRS matrix of genotype BLUPs

B mixed model with genotype and *subpopulation* or *cross* effects, resulting NIRS matrix of genotype BLUPs only

C: mixed model with genotype and *subpopulation* or *cross* effects, resulting NIRS matrix of genotype + *subpopulation* or *cross* BLUPs

All mixed models were fitted after der1 pre-process.

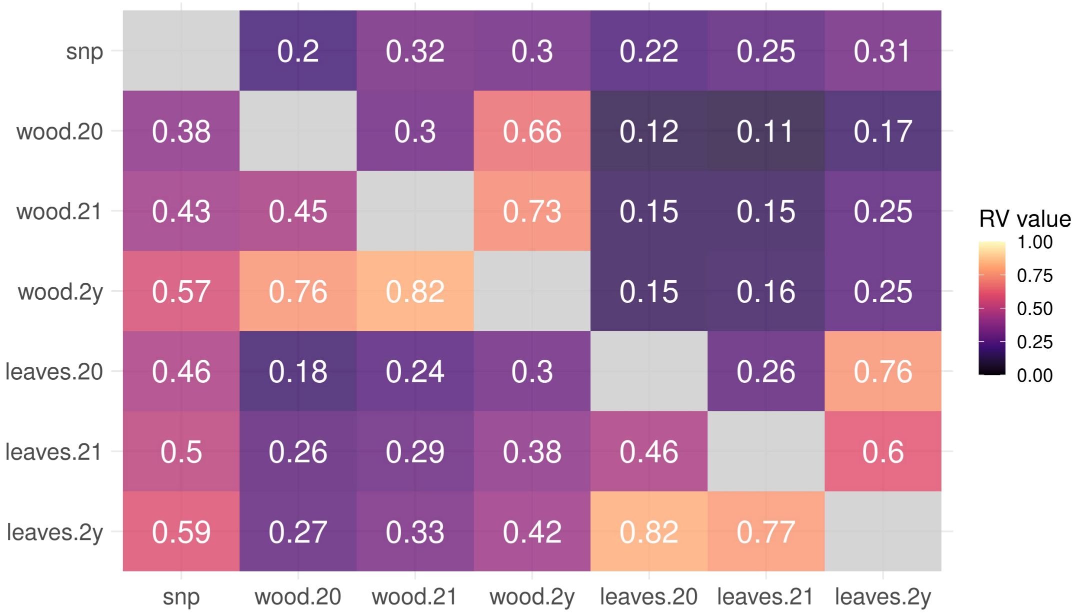


#### Figure S3 Co-inertia analysis between SNP, single and multi-year wood and leaves NIRS matrices

Upper triangle: in the diversity panel, lower triangle: in the half-diallel
Suffixes “20”, “21”, “2y” correspond to 2020, 2021 and multi-year NIRS, respectively. All mixed models were fitted after der1 pre-process.

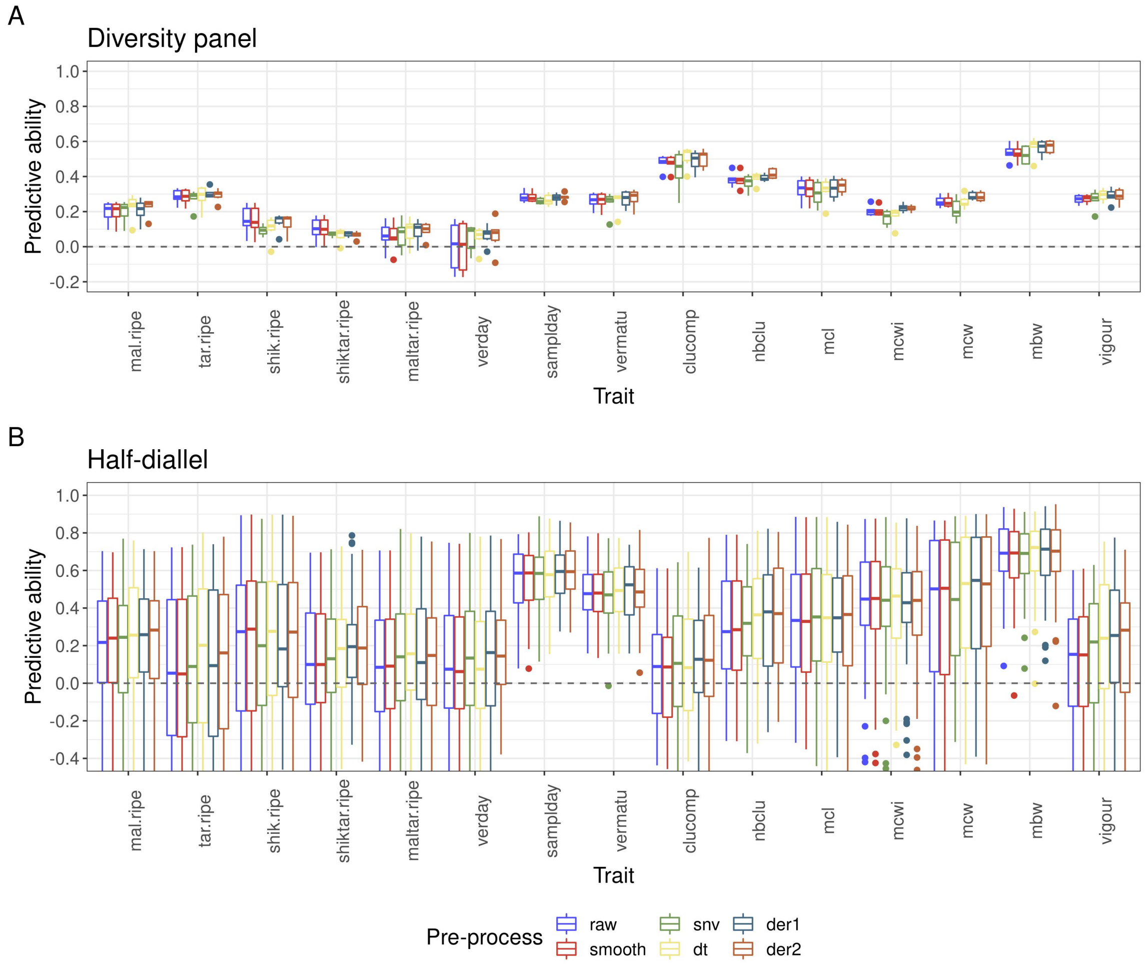


### Figure S4 Distribution of phenomic prediction predictive ability per pre-process and per trait

A: in the diversity panel with rrBLUP method (implemented with glmnet); B: in the half-diallel with HBLUP method (implemented with lme4GS).

Distribution of predictive ability is displayed across two tissue x three year BLUP models, times ten crosses in the half-diallel.

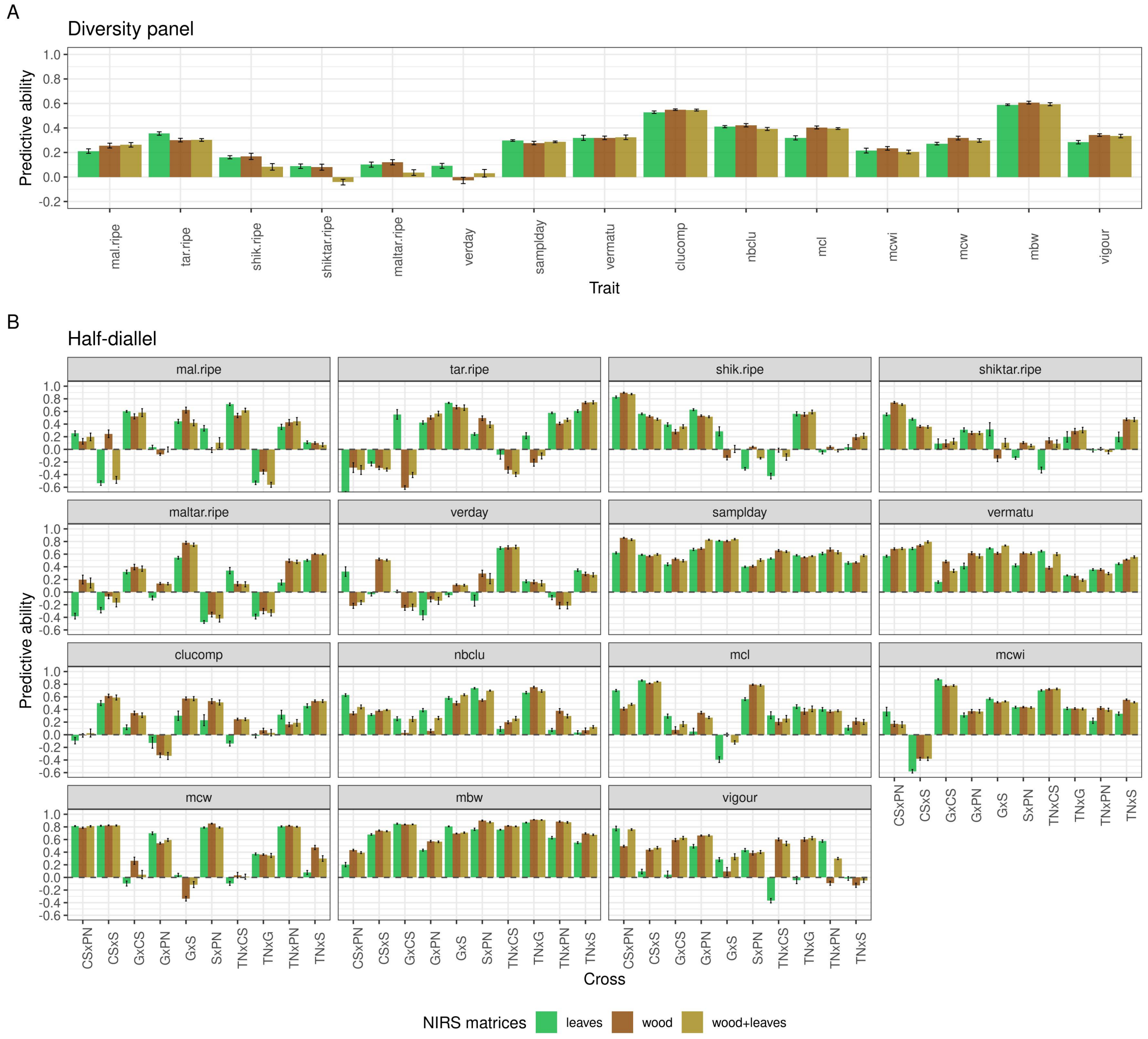


### Figure S5 Predictive ability of phenomic prediction with a single vs both tissues

For “2 years” NIRS BLUPs derived after der1 pre-process. A: in the diversity panel, B: in the half-diallel. Predictive ability values are displayed per trait for both populations, and also per cross in the half-diallel. Prediction models were fitted with glmnet in the diversity panel (except for wood+leaves configuration) and with lme4GS in the half-diallel. Error bars correspond to 95% confidence intervals around the mean, calculated for the ten CV repetitions.

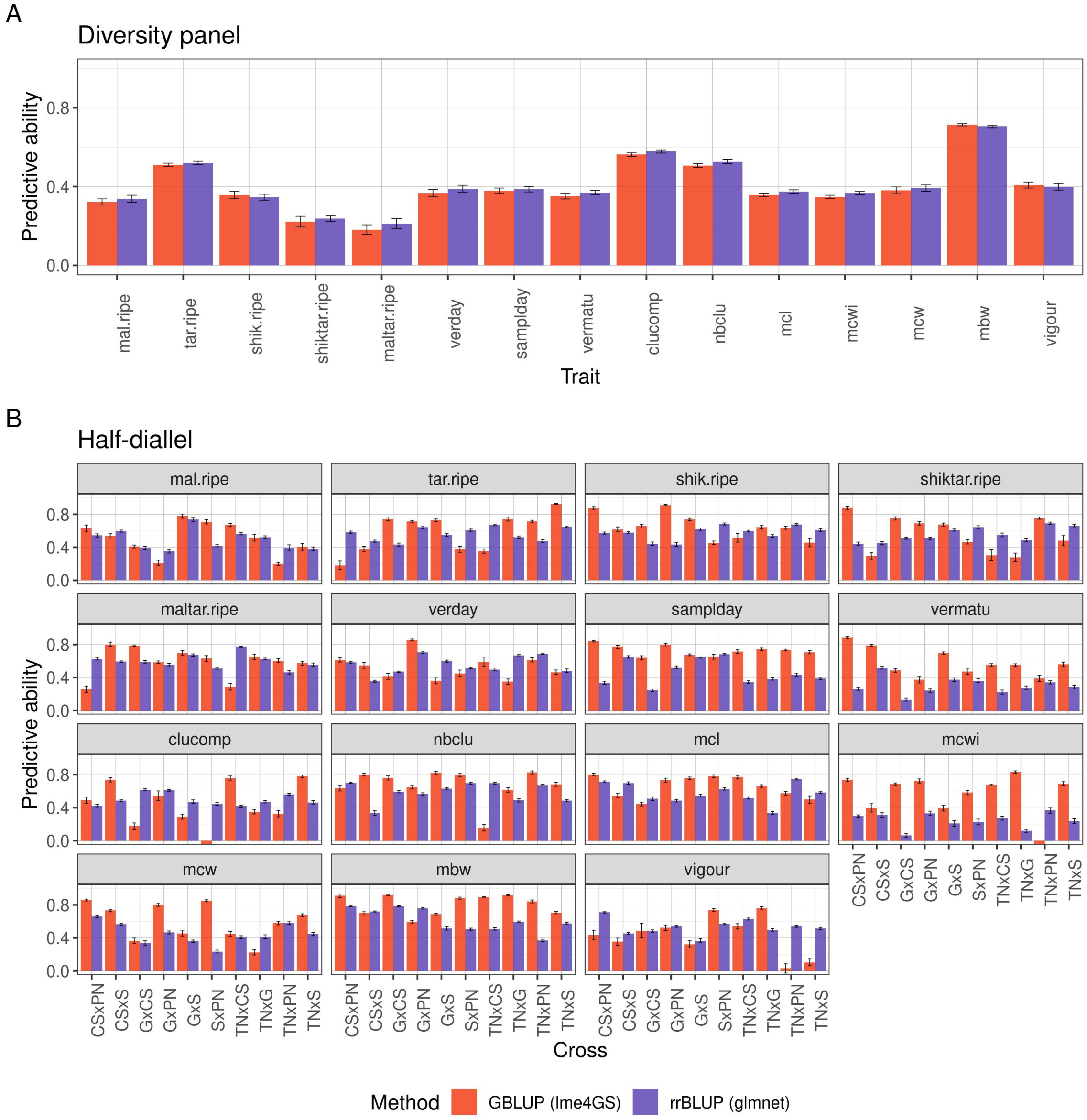


### Figure S6 Comparison of methods for genomic prediction

A: in the diversity panel; B: in the half-diallel. Predictive ability values are displayed per trait for both populations, and also per cross in the half-diallel. Error bars correspond to 95% confidence intervals around the mean, calculated for the ten CV repetitions.

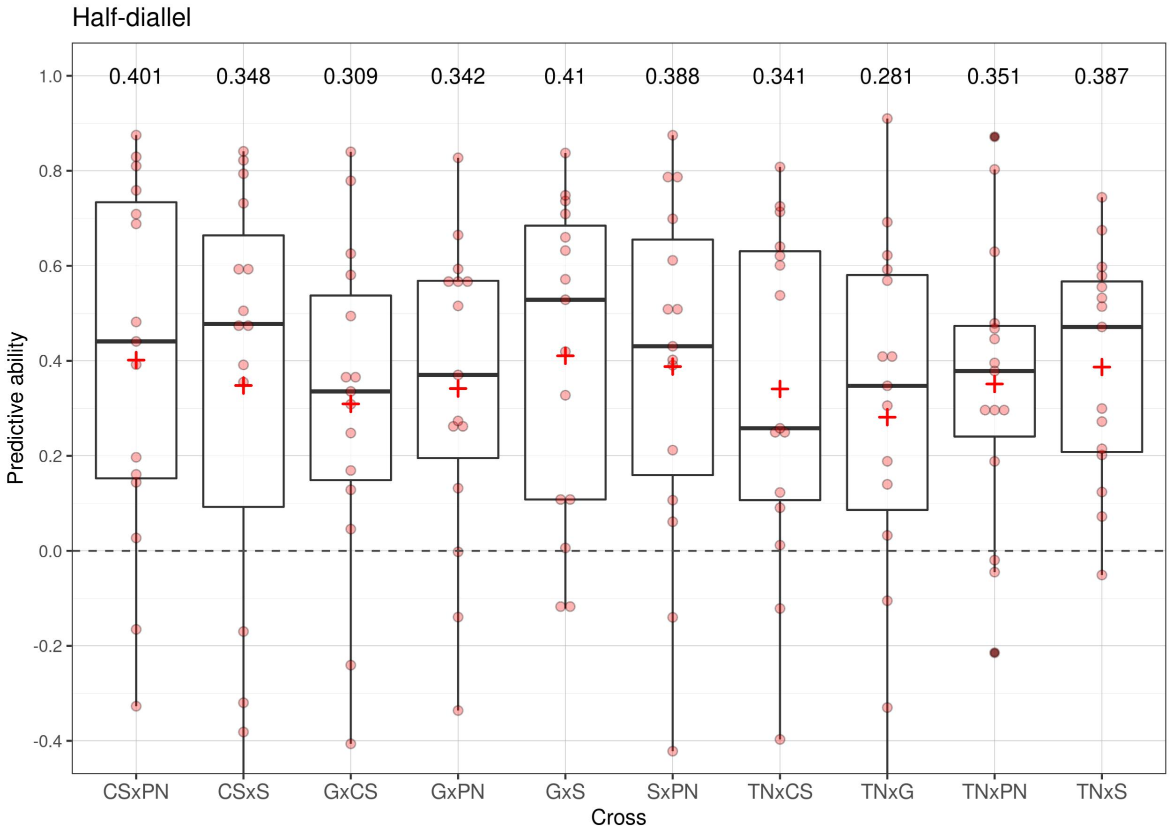


#### Figure S7 Distribution of phenomic prediction predictive ability over 15 traits in each half-diallel cross

For “2 years” NIRS BLUPs derived after der1 pre-process. Average PA per cross is displayed above each cross and with a red cross. Prediction models were fitted with lme4GS and included both wood and leaves NIRS relationship matrices.

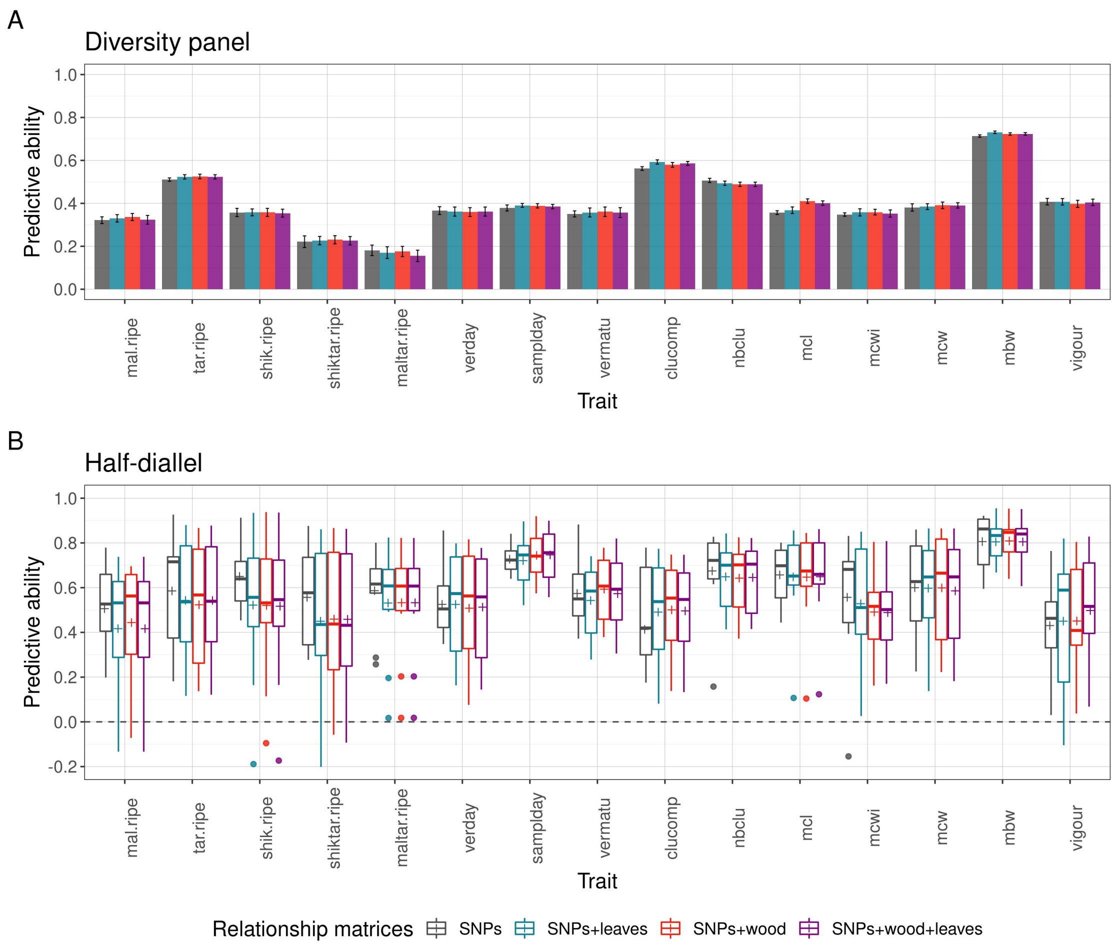


### Figure S8 Predictive ability of combined vs genomic prediction models

A: in the diversity panel (error bars correspond to 95% confidence intervals around the mean, calculated for the ten CV repetitions); B: in the half-diallel (distribution over 10 crosses and 10 CV repetitions). Prediction models were fitted with lme4GS and included as relationship matrices: SNPs, SNPs and leaves NIRS, SNPs and wood NIRS or SNPs, wood and leaves NIRS. Wood and leaves NIRS BLUPs were derived from a mixed model including both years, after der1 pre-process
